# Single-cell multiome of the human retina and deep learning nominate causal variants in complex eye diseases

**DOI:** 10.1101/2022.03.09.483684

**Authors:** Sean K. Wang, Surag Nair, Rui Li, Katerina Kraft, Anusri Pampari, Aman Patel, Joyce B. Kang, Christy Luong, Anshul Kundaje, Howard Y. Chang

## Abstract

Genome-wide association studies (GWAS) of eye disorders have identified hundreds of genetic variants associated with ocular disease. However, the vast majority of these variants are noncoding, making it challenging to interpret their function. Here, we present a joint single-cell atlas of gene expression and chromatin accessibility of the adult human retina with >50,000 cells, which we used to analyze noncoding single-nucleotide polymorphisms (SNPs) implicated by GWAS of age-related macular degeneration, glaucoma, diabetic retinopathy, myopia, and type 2 macular telangiectasia. We integrate this atlas with a HiChIP enhancer connectome, expression quantitative trait loci (eQTL) data, and base-resolution deep learning models to predict noncoding SNPs with causal roles in eye disease, assess SNP impact on transcription factor binding, and define their known and novel target genes. Our efforts nominate pathogenic SNP-target gene interactions for multiple vision disorders and provide a potentially powerful resource for interpreting noncoding variation in the eye.

## INTRODUCTION

Genome-wide association studies (GWAS) of eye disorders such as glaucoma, myopia, and age-related macular degeneration (AMD) have uncovered hundreds of genetic polymorphisms associated with ocular disease^1–5^. However, the vast majority of variants identified by GWAS reside in noncoding regions of the genome, making it challenging to interpret their function^6^. To better understand how noncoding variants mechanistically contribute to ocular pathology, it would be valuable to map in which cell types their corresponding loci are active. This information would provide novel insights into the cellular biology of genetically complex eye diseases and help nominate specific cell types as targets for therapies.

A recent advance in studying the noncoding genome has been the development of single-cell multiomic technologies such as paired single-cell RNA sequencing (scRNA-seq) and single-cell assay for transposase-accessible chromatin sequencing (scATAC-seq). While scRNA-seq can classify the different cell types of a tissue based on their transcriptional profiles, its combination with scATAC-seq allows for the additional mapping of cell type-specific chromatin accessibility. Together, these techniques can reveal the activity of noncoding DNA elements identified by GWAS and have been used to interrogate risk variants for conditions including Alzheimer’s disease, Parkinson’s disease, autism spectrum disorder, and autoimmunity ^7–9^.

Investigations into the noncoding genome have likewise benefitted from analytical innovations, such as the application of convolutional neural network (CNN)-based deep learning to predict the effects of noncoding polymorphisms^10–12^. Progress in this area has recently led to models with resolution down to a single nucleotide, enabling accurate determination of the critical bases within *cis*-regulatory sequences^8,12,13^. These models offer a validated approach to prioritize noncoding variants with functional relevance and are particularly suitable for tissues in which experimental manipulation is difficult.

Here, we generated a joint scRNA- and scATAC-seq atlas of the adult human retina composed of >50,000 cells and used it to analyze noncoding single-nucleotide polymorphisms (SNPs) implicated by GWAS of five eye diseases: AMD, glaucoma, diabetic retinopathy (DR), myopia, and type 2 macular telangiectasia (MacTel). Layering this atlas with a HiChIP enhancer connectome^14^, expression quantitative trait loci (eQTL) data^15^, and base-resolution deep learning models^12^, we then predicted noncoding SNPs with causal roles in eye disease. Our efforts nominate pathogenic SNP-target gene interactions for multiple vision disorders and provide a potentially powerful resource for interpreting noncoding variation in the eye.

## RESULTS

### Single-cell multiomics reveal the gene expression and chromatin accessibility landscapes of cell types in the human retina

To generate a single-cell multiome of the human retina, we performed joint scRNA- and scATAC-seq profiling on eight postmortem retinas from four individuals who had no history of eye disease (Supplementary Table 1). Following quality control filtering (Extended Data Fig. 1a-f) and removal of putative doublets (Extended Data Fig. 2a,b), we obtained a total of 51,645 human retinal cells in 22 clusters which we assigned to 13 different cell types (Fig. 1a,b and Extended Data Fig. 2c). These included abundant cell types like rod photoreceptors and Müller glia, as well as rarer cell types such as astrocytes and microglia, which each constituted only 0.4% of profiled cells (Fig. 1c and Supplementary Table 2). Consistent with published scRNA-seq studies of the human retina^16–18^, we observed cell type-specific expression of many genes, including *PDE6A* in rod photoreceptors, *GRIK1* in OFF-cone bipolar cells, *RLBP1* in Müller glia, *GRM6* in ON-cone and rod bipolar cells, *PRKCA* in rod bipolar cells, *ARR3* in cone photoreceptors, *GAD1* in GABAergic (GABA-) amacrine cells, *ONECUT1* in horizontal cells, *SLC6A9* in AII- and other glycinergic (gly-) amacrine cells, *NEFL* in retinal ganglion cells, *GJD2* in AII-amacrine cells, *GFAP* in astrocytes, and *C1QA* in microglia (Fig. 1d). In addition, we identified a list of candidate marker genes based on differential expression for each of the 13 cell types (Supplementary Data 1).

**Figure 1.**
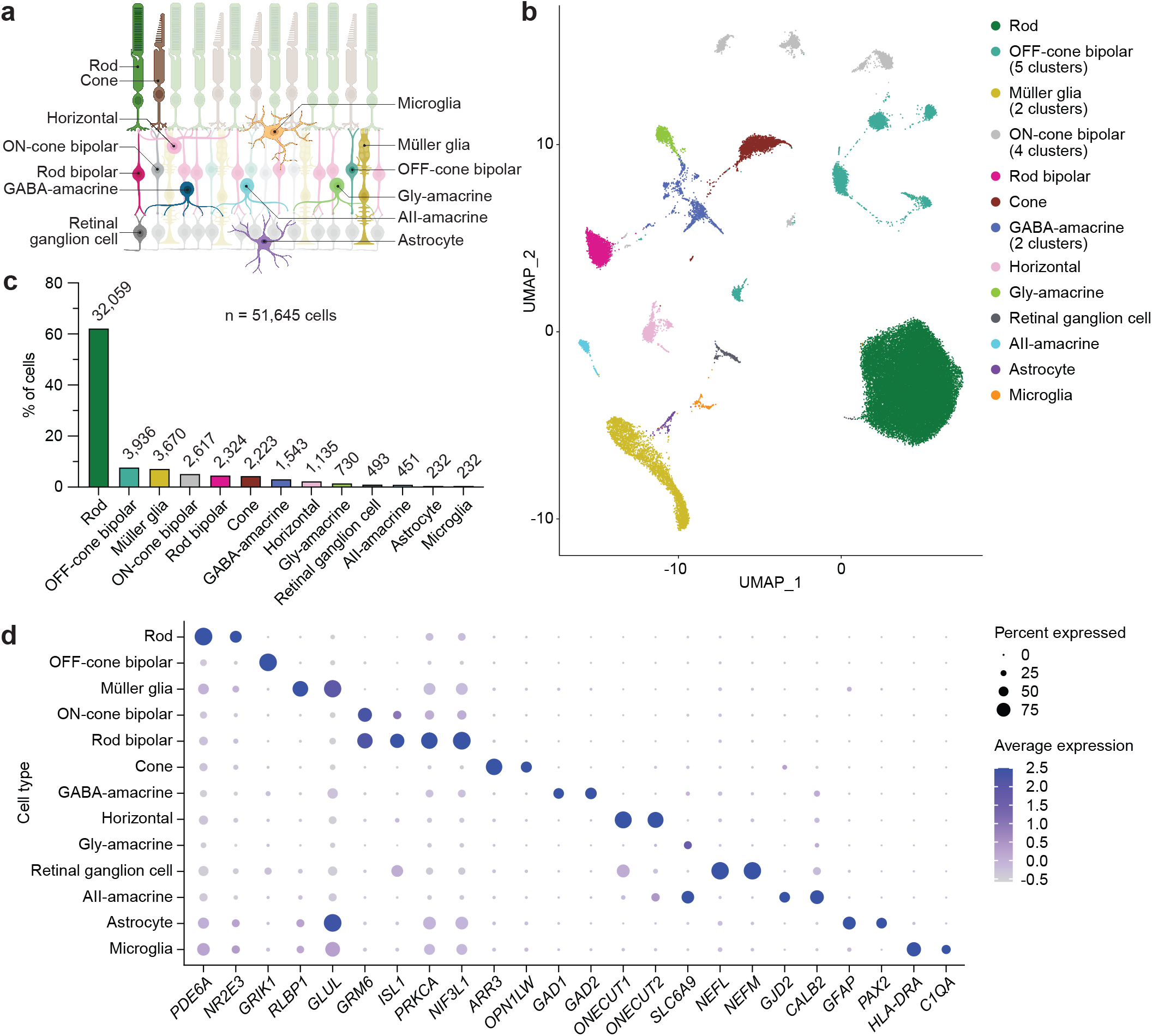
Transcriptional profiles from joint single-cell RNA- and ATAC-seq identify major cell types of the human retina. **a**, Schematic of the human retina depicting the cell types analyzed in this study. **b**, Uniform manifold approximation and projection (UMAP) plot of the 51,645 human retinal cells detected by scRNA-seq after quality control filtering and removal of putative doublets. Eight postmortem retinas from four donors were profiled. A total of 22 clusters were resolved and assigned to 13 cell types. **c**, Frequency of different cell types in the human retina as determined by scRNA-seq. Numbers above each bar denote absolute counts out of 51,645. **d**, Dot plot visualizing the normalized RNA expression of selected marker genes by cell type. The color and size of each dot correspond to the average expression level and fraction of expressing cells, respectively.

Using shared barcodes from joint multiomic profiling, we next assigned scATAC-seq profiles to the 13 cell types characterized above by scRNA-seq. Peak calling performed on scATAC-seq profiles from each cell type combined into pseudo-bulk ATAC replicates uncovered a total of 620,386 chromatin accessibility peaks (Fig. 2a and Supplementary Data 2). These scATAC peaks included >90% of peaks from published bulk ATAC-seq of the human retina (Fig. 2b)^19^, indicating that single-cell multiomics can recapitulate bulk ATAC-seq data. Conversely, more than half of scATAC peaks were unique to the single-cell dataset (Fig. 2b), and nearly 40% of scATAC peaks were accessible in only one cell type (Extended Data Fig. 3a). Supporting this, we found 197,826 scATAC marker peaks enriched in a cell type-specific manner (Fig. 2c and Supplementary Data 3), including many located near cell type-specific genes (Fig. 2d).

**Figure 2.**
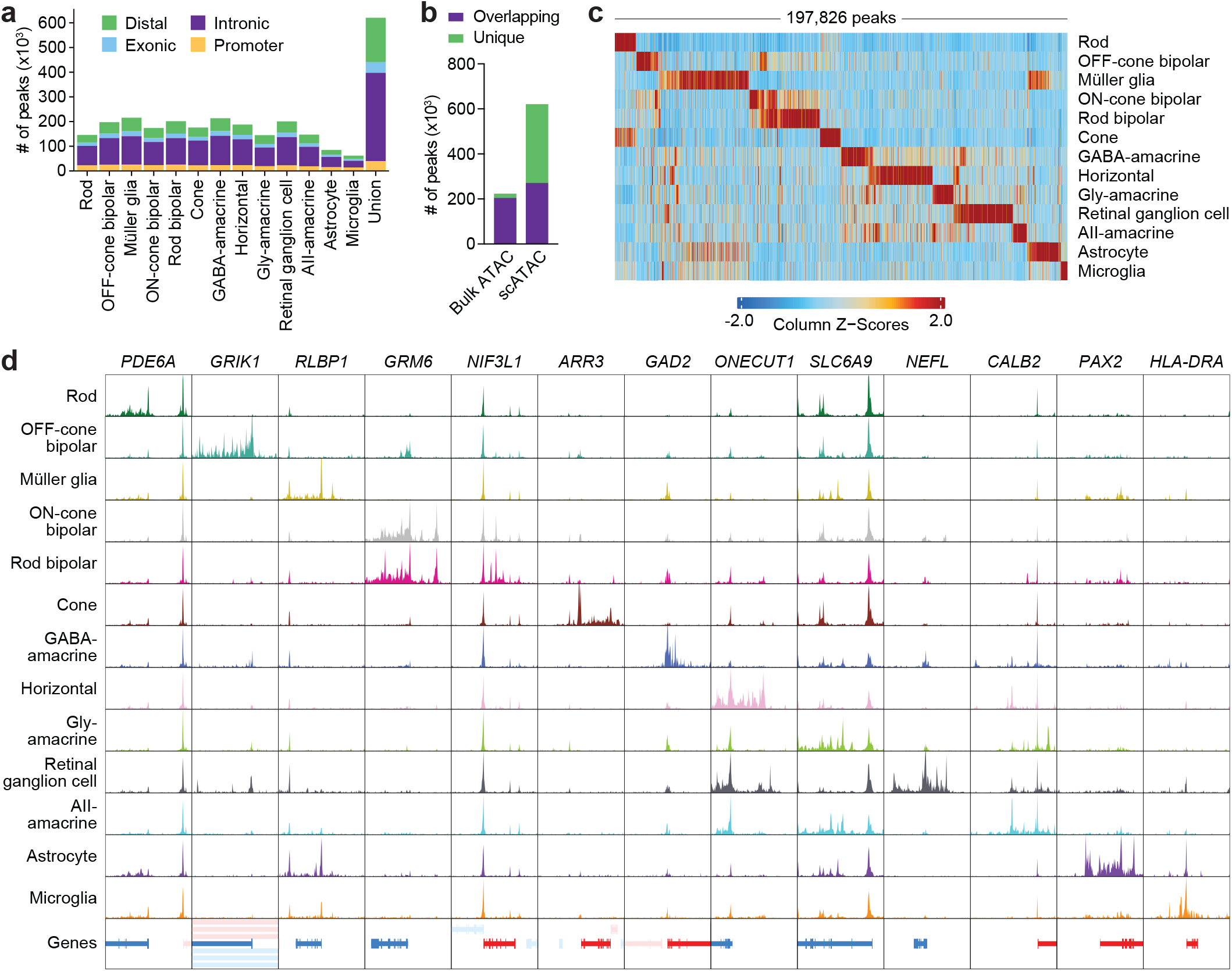
Chromatin accessibility profiles from joint single-cell RNA- and ATAC-seq of the human retina reveal cell type-specific epigenetic landscapes. **a**, Number of chromatin accessibility peaks for each cell type as determined by scATAC-seq. Peaks were required to be present in a least two pseudo-bulk ATAC replicates (n = 2 for Astrocyte and Microglia, n = 5 for all other cell types). **b**, Overlap of scATAC peaks with peaks from published human retina bulk ATAC-seq data. Overlapping was defined as peaks with any overlapping bases. **c**, Heatmap of scATAC marker peaks enriched in each cell type. Each column represents a marker peak. **d**, Sequencing tracks of chromatin accessibility near selected marker genes by cell type. Each track represents the aggregate scATAC signal of all cells from the given cell type normalized by the total number of reads in TSS regions. Genes in the sense direction (TSS on the left) are shown in red, while genes in the antisense direction (TSS on the right) are shown in blue. Coordinates for each region: *PDE6A* (chr5:149924792-149964793), *GRIK1* (chr21:29905031-29955033), *RLBP1* (chr15:89201750-89241751), *GRM6* (chr5:178975297-179015298), *NIF3L1* (chr2:200874325-200914327), *ARR3* (chrX:70248304-70288305), *GAD2* (chr10:26186306-26246307), *ONECUT1* (chr15:52781076-52821078), *SLC6A9* (chr1:44005465-44035467), *NEFL* (chr8:24937109-24977110), *CALB2* (chr16:71323711-71368713), *PAX2* (chr10:100715602-100755603), *HLA-DRA* (chr6:32419841-32459842).

With these scATAC peaks, we then conducted motif enrichment analysis to predict what transcription factors (TFs) might be active in each cell type (Fig. 3a and Supplementary Data 4). In accord with published literature, we observed enrichment of binding motifs for TFs with known cell type-specific functions, such as OTX2 in photoreceptors and bipolar cells^20^, ONECUT family members in horizontal cells^21^, POU4F family members in retinal ganglion cells^22^, and SPI1 (PU.1) in microglia^23^. For some TFs, cell type-specific activity was also supported by footprinting analysis of scATAC peaks (Fig. 3b), which revealed motif centers to be protected from Tn5 transposition, consistent with TF occupancy. These data offer a cell typespecific catalog of candidate TFs in the adult retina and may aid our understanding of gene-regulatory networks controlling vision.

**Figure 3.**
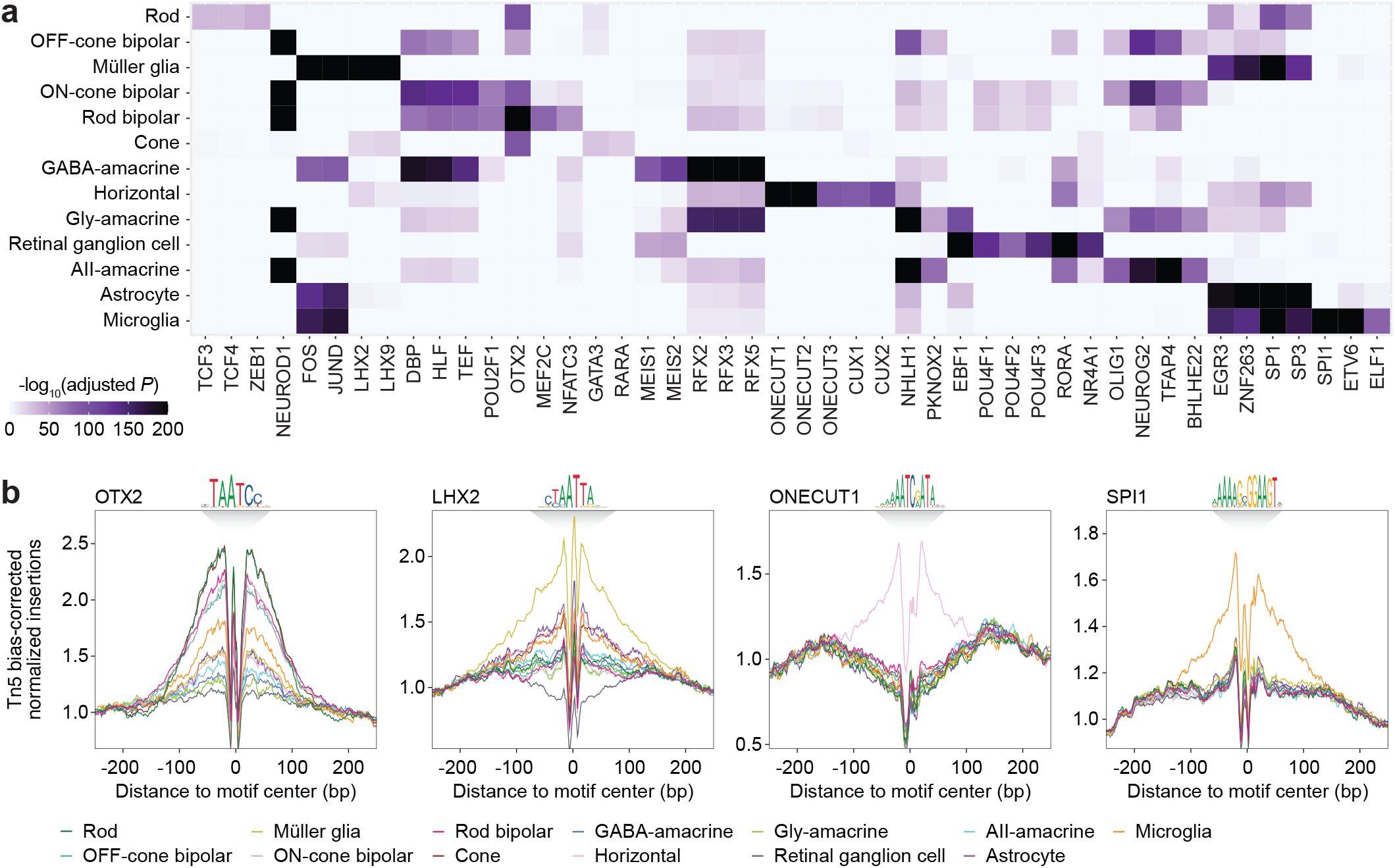
Motif analysis of accessible DNA regions in the human retina predicts cell typespecific transcription factors. **a**, Heatmap of selected TF binding motifs enriched in each cell type. Darker colors indicate more significant enrichment. **b**, Footprinting analysis of selected TFs across cell types. Footprints were corrected for Tn5 insertion bias by dividing the footprinting signal by the Tn5 insertion signal.

### Single-cell multiomics uncover the cellular contexts of variants implicated by ocular disease GWAS

Using our single-cell multiome, we sought to better understand risk loci identified by GWAS of complex eye disorders. To this end, we compiled of a list of 1,331 unique index SNPs from the NHGRI-EBI GWAS Catalog representing GWAS hits for five eye diseases: AMD, glaucoma, DR, myopia, and MacTel (Fig. 4a and Supplementary Table 3)^24^. The vast majority (96.5%) of these SNPs localize to noncoding regions of the genome and thus cannot be interpreted with scRNA-seq data alone. We performed linkage disequilibrium (LD) expansion on all index SNPs to include nearby variants with high probability of coinheritance (LD R^2^ >0.9 based on phase 3 genotypes from the 1000 Genomes Project) (Extended Fig. 3b)^25^. From this, we obtained a total of 7,034 unique noncoding SNPs in loci associated with eye disorders (Supplementary Data 5).

**Figure 4.**
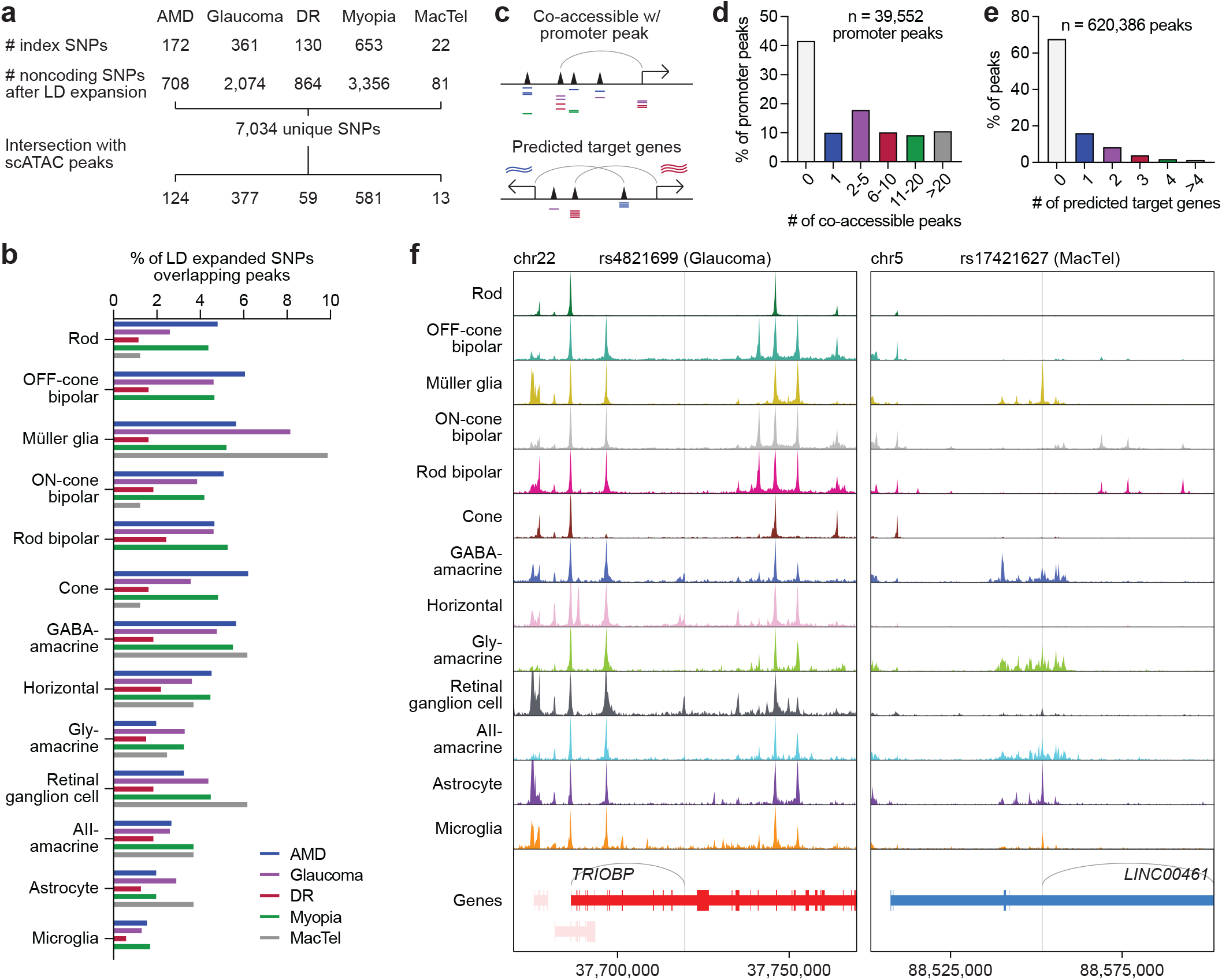
Single-cell multiomics pinpoint the cellular targets of noncoding variants in eye diseases. **a**, Overview of SNP selection for interrogating ocular disease GWAS. Index SNPs obtained from GWAS of each disease were subjected to LD expansion, and the resulting noncoding SNPs intersected with scATAC peaks. **b**, Percentage of LD expanded SNPs from each disease that overlapped with chromatin accessibility peaks for each cell type. **c**, Schematic of promoter coaccessibility and predicted target gene analyses. Co-accessible was defined as scATAC peaks whose accessibility showed a correlation score >0.3. Predicted target genes were defined as genes whose RNA expression showed a correlation score >0.3 relative to the accessibility of the tested scATAC peak. **d**, Number of scATAC peaks co-accessible with each promoter peak. **e**, Number of predicted target genes for each scATAC peak. **f**, Sequencing tracks of chromatin accessibility near rs4821699 (chr22:37719685) and rs17421627 (chr5:88551768). Genes in the sense and antisense directions are shown in red and blue, respectively. The location of each SNP is depicted by a vertical gray line. Gray arcs indicate predicted target genes for the scATAC peak containing the SNP of interest.

To determine in which retinal cell types each of the 7,034 SNPs might be active, we overlapped SNP locations with scATAC peaks from our dataset. We found that 1,152 SNPs (16.4%) overlapped with a scATAC peak (Fig. 4b and Extended Data Fig. 3c), and that most SNP-containing peaks were present in only one or two cell types (Extended Fig. 3d). We next conducted two orthogonal analyses to refine our list of SNPs for those more likely to possess gene regulatory functions (Fig. 4c). First, we identified SNPs in scATAC peaks that were co-accessible with peaks in promoter regions, reasoning that this would select for SNPs in active enhancers. We found 39,552 such promoter peaks in the human retina, 58.3% of which were co-accessible with at least one scATAC peak (Fig. 4d). Leveraging our paired scRNA- and scATAC-seq data, we also searched for SNPs in peaks that had at least one predicted target gene based on same-cell correlations between peak accessibility and gene expression. Using this method, we predicted target genes for 199,055 (32.1%) of the 620,386 scATAC peaks in our dataset (Fig. 4e). For nearly half (44.3%) of these peaks, our predictions differed from the nearest gene on the linear genome (Extended Data Fig. 3e), suggesting that noncoding SNPs do not necessarily regulate their nearest gene.

We identified 241 SNPs in scATAC peaks that were co-accessible with promoter peaks and 374 SNPs that had predicted target genes, with 202 SNPs meeting both criteria. As an example, we examined rs4821699 residing in an intron of *TRIOBP* on chromosome 22. This locus has been implicated in glaucoma by multiple GWAS and encodes a protein thought to regulate cytoskeletal organization^2,26,27^. We observed that rs4821699 was most accessible in retinal ganglion cells (Fig. 4f), the major cell type that undergoes degeneration during glaucoma. Based on correlations with gene expression, the peak containing this SNP was furthermore predicted to target *TRIOBP*. We hypothesize that rs4821699 might therefore play a role in glaucoma by altering *TRIOBP* expression in retinal ganglion cells.

A handful of SNPs associated with eye diseases have been experimentally studied using retinal organoids derived from induced pluripotent stem cells. One such SNP is rs17421627, an index SNP from GWAS of MacTel representing a T-to-G substitution on chromosome 5^5,28^. We determined rs17421627 to be one of only five SNPs for MacTel with a predicted target gene and found the SNP to be most accessible in Müller glia and astrocytes (Fig. 4f). Using linked gene expression data, we also predicted rs17421627 to act on *LINC00461,* a long noncoding RNA. Consistent with these predictions, deletion of the locus containing rs17421627 in human retinal organoids has been shown to significantly downregulate *LINC00461* with the strongest effect in Müller glia^29^. These examples illustrate how single-cell multiomics can reveal the cellular targets of noncoding variants in the retina and nominate how they might contribute to eye disorders.

### Integration of single-cell multiome with HiChIP and eQTL data validates SNP-target gene predictions

To further prioritize our list of SNPs, we combined our data with two complementary methods for identifying functional SNP-gene interactions genome-wide (Fig. 5a). We first performed HiChIP for acetylated histone H3 lysine 27 (H3K27ac), a mark of active enhancers and promoters^30^, to characterize the three-dimensional (3D) enhancer “connectome” of the human retina (Extended Data Fig. 4a,b)^14,31^. We uncovered 16,692 loop anchors connected by 9,670 HiChIP loops, including several linking regions of chromatin accessibility to the transcription start sites (TSSs) of cell type-specific genes (Extended Data Fig. 4c). Of these loops, >95% overlapped with a scATAC peak in both anchors, and >99% overlapped with a peak in at least one anchor (Fig. 5b). This result shows that accessible chromatin sites identified in scATAC-seq data possess biochemical characteristics of active enhancers and supports their connection to target genes. We additionally analyzed our list of SNPs using published human retina eQTL data from the Eye Genotype Expression (EyeGEx) database^15^. For >90% of SNPs in scATAC peaks, retina eQTL data was available (Fig. 5c), enabling genes whose mRNA expression in the human retina changed with specific SNPs to be identified at the bulk tissue level.

**Figure 5.**
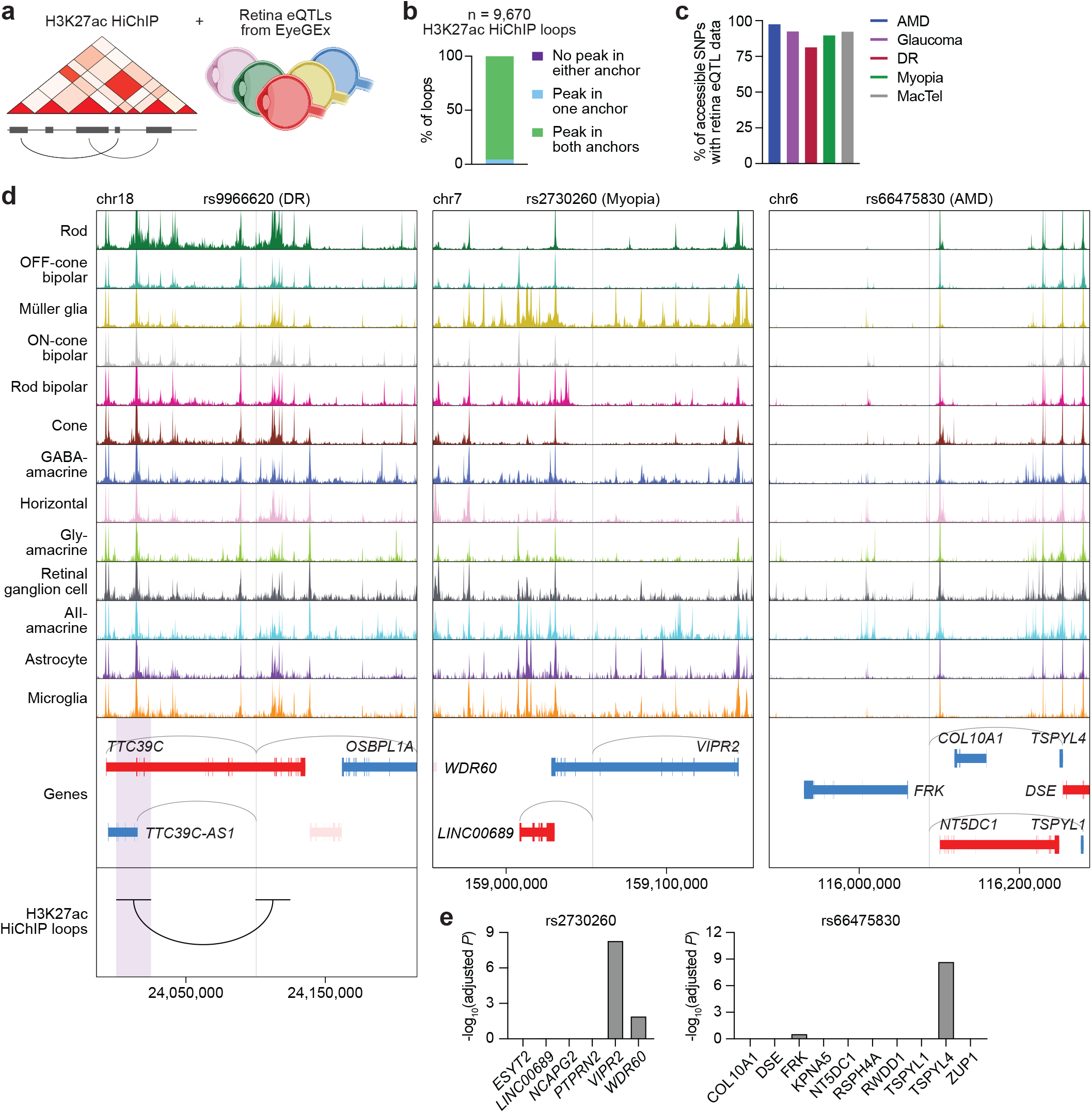
Integration of single-cell multiome with HiChIP and eQTL data prioritizes functional noncoding polymorphisms in the human retina. **a**, Schematic of H3K27ac HiChIP and eQTL analyses used to prioritize SNPs. **b**, Overlap of HiChIP loop anchors (n = 2 biological replicates) with scATAC peaks. **c**, Percentage of SNPs in scATAC peaks for each disease with available retina eQTL data. **d**, Sequencing tracks of chromatin accessibility near rs9966620 (chr18:24100771), rs2730260 (chr7:159054238), and rs66475830 (chr6:116087639). Genes in the sense and antisense directions are shown in red and blue, respectively. The location of each SNP is depicted by a vertical gray line. Gray arcs indicate predicted target genes for the scATAC peak containing the SNP of interest. The black arc overlapping with rs9966620 indicates a H3K27ac HiChIP loop with the region encompassed by the opposite anchor highlighted in purple. **e**, Significance of SNP-gene associations for rs2730260 or rs66475830 and their nearby genes as determined by retina eQTL analysis. Adjusted *P* values for each gene were calculated by multiplying the nominal *P* value listed in the EyeGEx database by the number of SNP-gene pairs tested for that SNP.

We found 187 disease-associated SNPs in scATAC peaks that were linked to a gene by a H3K27ac HiChIP loop. These included rs9966620, the top SNP from a GWAS of DR representing a G-to-A transition in an intron of *TTC39C* on chromosome 18^32^. Using our multiome, we determined that the scATAC peak containing rs9966620 was most accessible in rods (Fig. 5d). However, this peak also correlated with the expression of multiple target genes, hampering efforts to interpret how the SNP might function. Incorporating our HiChIP data, we were able to locate a 3D loop connecting rs9966620 with a region 75 kilobases (kb) upstream. This region intersected the TSS of only one gene, *TTC39C-AS1,* suggesting that rs9966620 may modulate DR risk by interacting with *TTC39C-AS1* in rods.

We additionally detected 596 disease-associated SNPs in scATAC peaks that were significantly associated with a gene by eQTL analysis. One example was rs2730260, a SNP in an intron of *VIPR2* that has been implicated in myopia^33^. This locus encodes one of two known receptors for vasoactive intestinal peptide (VIP), a signaling molecule involved in visual processing^34^. We found that rs2730260 resided in a chromatin accessibility peak specific to Müller glia that again had multiple predicted target genes (Fig. 5d). This ambiguity was clarified by retina eQTL data, which showed that variation at rs2730260 significantly correlated with the expression of only *VIPR2* (Fig. 5e), supporting this gene as the SNP’s primary target. Integration of eQTL data similarly improved our interpretation of rs66475830 on chromosome 6 in the *FRK-NT5DC1-COL10A1* risk locus for AMD^35,36^. This region contains nearly 20 genes within a span of a megabase (Mb), making it particularly difficult to functionally annotate GWAS hits. From our single-cell data, we determined rs66475830 to be accessible in amacrine and horizontal cells and predicted *TSPYL1* and *TSPYL4* as target genes (Fig. 5d). Retina eQTL analysis revealed that variation at this position was significantly associated with *TSPYL4* expression, but not that of other nearby genes (Fig. 5e), nominating *TSPYL4* as the effector gene of rs66475830.

Lastly, we identified many SNP-target gene relationships supported by both HiChIP and eQTL data, such as rs77272443 and rs4102217 located in risk loci for myopia and glaucoma, respectively^37,38^. For both of these SNPs, HiChIP and eQTL analyses again refined target gene predictions (Extended Data Fig. 5a,b), demonstrating how the combination of single-cell multiomics with other assays can enhance interpretation of noncoding variants in eye disease.

### Integration of single-cell multiome with base-resolution deep learning nominates functional mechanisms for disease-associated SNPs

CNN-based deep learning models have proven capable of discerning disease-associated SNPs from other noncoding variants^8,10,11^. As a final method to prioritize SNPs in our dataset, we therefore trained CNNs derived from the BPNet architecture on scATAC-seq profiles for each of the 13 retinal cell types (Fig. 6a and Extended Data Fig. 6a,b)^12^. At each SNP region, we compared the projected per-base change in chromatin accessibility between reference and alternate alleles using models specific to the different cell types. These calculations allowed us to identify “high effect” SNPs, which we defined as SNPs predicted to cause a statistically significant (false discovery rate <0.01) absolute log2 fold change of allele-specific read counts >0.5 in local chromatin accessibility in any cell type.

**Figure 6.**
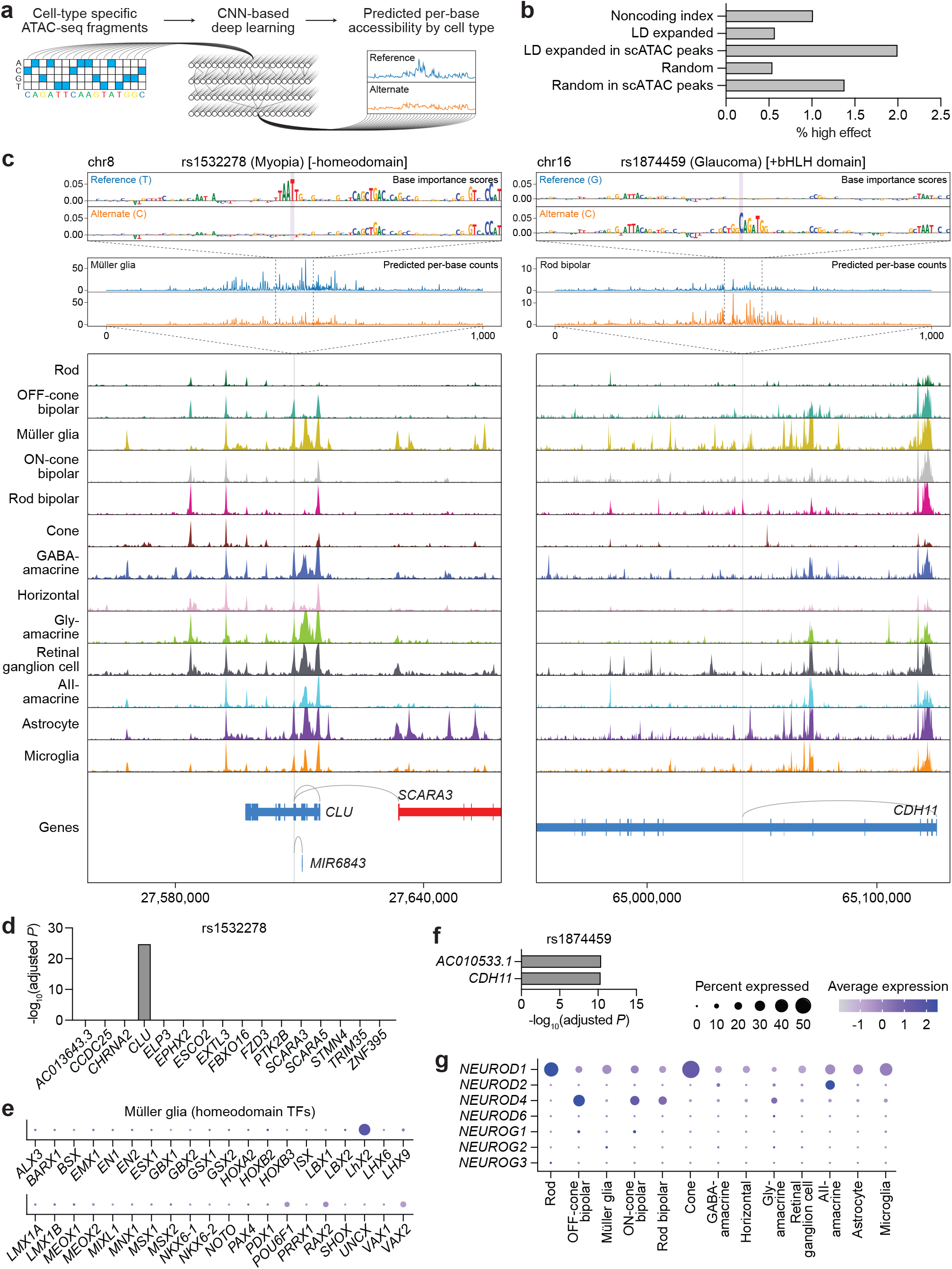
Integration of single-cell multiome with base-resolution deep learning nominates functional mechanisms for disease-associated SNPs. **a**, Schematic of the CNN-based deep learning pipeline. **b**, Percentage of noncoding index SNPs (n = 1,284), LD expanded SNPs (n = 7,034), LD expanded SNPs in scATAC peaks (n = 1,152), randomly selected GC-matched SNPs (n = 9,984), and randomly selected SNPs in scATAC peaks (n = 1,160) that were categorized as high effect. **c**, Top: Predicted per-base accessibility for rs1532278 (chr8:27608798) and rs1874459 (chr16:65041801) in Müller glia and rod bipolar cells, respectively, as determined by deep learning models. A 100-bp window depicts the importance of each base to predicted accessibility at the SNP, and a 1,000-bp window depicts predicted per-base counts for the reference (blue) and alternate (orange) alleles. SNP bases are highlighted in purple. For rs1874459, similar changes in accessibility were predicted for OFF-cone bipolar, ON-cone bipolar, gly-amacrine, and AII-amacrine cells. Bottom: Sequencing tracks of chromatin accessibility near rs1532278 and rs1874459. Genes in the sense and antisense directions are shown in red and blue, respectively. The location of each SNP is depicted by a vertical gray line. Gray arcs indicate predicted target genes for the scATAC peak containing the SNP of interest. **d,f**, Significance of SNP-gene associations for rs1532278 (d) or rs1874459 (f) and their nearby genes as determined by retina eQTL analysis. Adjusted *P* values for each gene were calculated by multiplying the nominal *P* value listed in the EyeGEx database by the number of SNP-gene pairs tested for that SNP. **e**, Dot plot visualizing the normalized RNA expression of 40 different homeodomain TFs in Müller glia. The selected TFs correspond to the 40 homeodomain factors whose binding motifs were most significantly enriched in Müller glia as determined by motif analysis (Supplementary Data 4). **g**, Dot plot visualizing the normalized RNA expression of neuroD and neurogenin family members by cell type.

We found 23 SNPs (2.0%) residing in scATAC peaks that qualified as high effect, a greater percentage than among index SNPs, LD expanded SNPs, random SNPs matched for GC content, and random SNPs residing in scATAC peaks (Fig. 6b and Supplementary Data 7). One of the top scoring SNPs was rs1532278, an index SNP associated with myopia and residing in an intron of *CLU* on chromosome 8^3^. Our atlas predicted rs1532278 to regulate *CLU,* a notion reinforced by eQTL data, and determined the SNP to be accessible in nine of 13 retinal cell types (Fig. 6c,d). Despite this, base-resolution models projected a T-to-C transition at rs1532278 to alter chromatin accessibility only in Müller glia, specifically by disrupting the motif of a homeodomain TF. Our findings suggest that even though rs1532278 is accessible across multiple cell types, its functional impact in the retina might be restricted to Müller glia due to a cell type-specific homeodomain TF. We speculate that this TF could be LHX2 given its robust expression in Müller glia by both our scRNA-seq data (Fig. 6e) as well as data from animal models^39^.

Another high effect SNP was rs1874459 located in an intron of *CDH11* on chromosome 16, a locus implicated by multiple GWAS for glaucoma^2,26^. Using our multiome, we found rs1874459 to be most accessible in rod bipolar cells and predicted *CDH11* as one of its target genes, an idea supported by eQTL data (Fig. 6c,f). Incorporating base-resolution models, we then determined that the G-to-C transversion represented by rs1874459 introduced a new basic helix-loop-helix (bHLH) domain, which was expected to increase accessibility in rod bipolar, OFF-cone bipolar, ON-cone bipolar, gly-amacrine, and AII-amacrine cells. Of the bHLH TFs, members of the neuroD and neurogenin families in particular were predicted by motif analysis to be significantly enriched in these five cell types (Fig. 3a and Supplementary Data 4). We thus compared all neuroD and neurogenin family members using our scRNA-seq data, which revealed only NEUROD4 to be specific to bipolar and amacrine cells (Fig. 6g), consistent with its role in specifying these cell types during development^40,41^. Together, our results suggest that rs1874459 may act on *CDH11* in bipolar and amacrine cells by creating a new bHLH domain recognized by NEUROD4.

## DISCUSSION

In this study, we applied single-cell multiomics, HiChIP, eQTL analysis, and base-resolution deep learning to the human retina to decipher the role of noncoding risk variants in five eye diseases. Integrating these methods allowed us to predict gene and cellular targets in the retina for hundreds of SNPs and nominate dozens as pathogenic and meriting functional validation. From an initial list of >7,000 noncoding SNPs, we identified 1,152 located in chromatin accessibility peaks. We subsequently focused on SNPs 1) that were co-accessible with a promoter, 2) whose accessibility correlated with the expression of a nearby gene, 3) that were linked to a gene in 3D space by a H3K27ac HiChIP loop, 4) that demonstrated significant association with a gene based on retina eQTL data, and 5) that were predicted to alter local chromatin accessibility as determined by base-resolution models. We propose that SNPs meeting most or all of these criteria (Extended Data Fig. 7a-e and Supplementary Data 5) be prioritized in future validation efforts.

Our findings build upon recent works that used primarily fetal tissue and stem cell-derived organoids to map cell type-specific chromatin accessibility in the human retina^29,42^. Datasets from these studies offer a rich resource for decoding retinal development, but might not fully recapitulate the biology of the mature retina, making them potentially less suitable for studying eye disorders that present later in life. Here, we not only generated a single-cell multiome of the adult human retina to pinpoint cellular targets for disease-associated SNPs, but also combined it with multiple orthogonal analyses to define putative SNP-target gene interactions. By performing base-resolution deep learning, we were further able to uncover insights not readily apparent from single-cell, HiChIP, and eQTL data, such as the predicted impact of SNPs on TF binding and the directionality of these effects. To facilitate its use, our atlas is publicly available at https://eyemultiome.su.domains/.

Finally, it should be noted that the majority of SNPs we examined did not overlap with any chromatin accessibility peaks, suggesting that they were not active in the retina. We hypothesize that many of these unassigned SNPs may instead function in other parts of the eye and thus could not be captured by our analysis. For instance, although the neural retina is damaged in AMD and DR, the retinal pigment epithelium and vasculature, respectively, are thought to be the primary sites of pathology^43,44^. In glaucoma, the trabecular meshwork and ciliary body can modulate disease severity as evidenced by treatments that act on these tissues^45^. Likewise, the choroid and sclera may be involved in myopia given that they elongate alongside the retina with increasing nearsightedness^46^. Multiomic characterization of these additional ocular regions would enable a more complete understanding of how noncoding SNPs contribute to vision disorders.

## METHODS

### Human tissues

Postmortem adult human retinas were procured from consented donors by Lions VisionGift (Portland, OR, USA) or Lions Gift of Sight (St Paul, MN, USA) under protocols approved by the Eye Bank Association of America. None of the donors had a history of ocular disease. Deidentified retinas were flash-frozen in liquid nitrogen with a maximum death-to-preservation interval of 12 hours and shipped to Stanford University for processing.

### Nuclei isolation

Nuclei were isolated from frozen retinas using the Omni-ATAC protocol (https://doi.org/10.17504/protocols.io.6t8herw)^47^. Briefly, tissues were Dounce homogenized in cold homogenization buffer containing 0.3% IGEPAL CA-630 in the presence of protease and RNase inhibitors to release nuclei from frozen cells. Nuclei were subsequently purified via iodixanol gradient centrifugation and washed with ATAC resuspension buffer containing RNase inhibitor and 0.1% Tween-20 before permeabilization following the 10x Genomics demonstrated protocol for complex tissues (CG000375, Rev. B). After resuspension in diluted nuclei buffer, nuclei were counted using a manual hemocytometer to achieve a targeted nuclei recovery of 10,000 nuclei per sample.

### scRNA- and scATAC-seq library generation

Joint scRNA- and scATAC-seq libraries were prepared using the 10x Genomics Single Cell Multiome ATAC + Gene Expression kit according to manufacturer’s instructions. Libraries were sequenced with paired-end 150-bp reads on an Illumina NovaSeq 6000 to a target depth of 250 million read pairs per sample.

### scRNA- and scATAC-seq data preprocessing and quality control

Demultiplexed scRNA- and scATAC-seq fastq files were inputted into the Cell Ranger ARC pipeline (version 2.0.0) from 10x Genomics to generate barcoded count matrices of gene expression and ATAC data. For each sample, count matrices were loaded in ArchR and selected for barcodes that appeared in both the scRNA-seq and scATAC-seq datasets^48^. Samples in ArchR were quality control filtered for nuclei with 200-50,000 RNA transcripts, <1% mitochondrial reads, <5% ribosomal reads, TSS enrichment >6, and >2,500 ATAC fragments. Quality control filtered nuclei subsequently underwent automated removal of doublets using the filterDoublets function in ArchR, which identifies and removes the nearest neighbors of simulated doublets^48^.

### scRNA-seq data analysis

scRNA-seq data from nuclei remaining after quality control filtering and automated removal of doublets were analyzed using Seurat (version 3.1.5)^49^. After merging all preprocessed samples into a single Seurat object, gene expression counts were normalized using the NormalizeData function, scaled using the ScaleData function, and batch corrected using Harmony^50^. Graphbased clustering was then performed on the Harmony-corrected data using the top 20 principal components at a resolution of 0.5. Cluster identities were manually annotated based on the expression of genes from published scRNA-seq studies of the human retina^16–18^. Marker genes for each cluster were additionally identified using the FindAllMarkers function with a minimum fraction of 0.5 and a log2 fold change of 1 (Supplementary Data 1). Clusters expressing canonical marker genes from different cell types were designated as putative doublets and excluded, after which re-clustering was performed using the same parameters. Clusters with no detected marker genes were also excluded, after which the dataset was also re-clustered. Clusters in the final dataset representing subpopulations of the same cell type were grouped together for downstream analyses.

### scATAC-seq data analysis

scATAC-seq data were analyzed using ArchR (version 1.0.1) based on barcoded cell type identities from scRNA-seq^48^. For each cell type, pseudo-bulk ATAC replicates were created using the addGroupCoverages function with default parameters, which generated between two to five replicates depending on how many cells of that type were present in each sample. Chromatin accessibility peaks on chromosomes 1-22 and X and outside of blacklist regions were then called using the addReproduciblePeakSet function and MACS2^51,52^, with scATAC peaks for each cell type defined as those present in at least two pseudo-bulk ATAC replicates (Supplementary Data 2). Marker peaks were identified using the getMarkerFeatures function with a log2 fold change ≥1 and false discovery rate ≤0.01 as determined by Wilcoxon pairwise comparisons (Supplementary Data 3). Promoter peaks were defined as scATAC peaks within 2,000 bp upstream or 100 bp downstream of a TSS, and peaks co-accessible to promoter peaks were identified using the getCoAccessibility function with a correlation cutoff of 0.3 and resolution of 1. Predicted target genes for each scATAC peak were generated using the getPeak2GeneLinks function integrating barcode-matched RNA expression data from scRNA-seq with a correlation cutoff of 0.3 and resolution of 1. Nearest genes were determined using the BEDTools closest function based on gene annotations from TxDb.Hsapiens.UCSC.hg38.knownGene^53^.

### Bulk ATAC-seq data analysis

Bulk ATAC-seq analysis was performed on published ATAC-seq data from five healthy human retinas^19^. After adapter trimming, fastq files were mapped to the hg38 genome using Bowtie2 and filtered to remove PCR duplicates and retain reads from only chromosomes 1-22 and X^54^. Peak calling was then conducted individually on each sample using MACS2^51^, followed by exclusion of peaks in blacklist regions^52^. Peak calls present in at least two of the five retinas were included in the bulk ATAC-seq peak set.

### Sequencing tracks

Sequencing tracks of chromatin accessibility were generated in ArchR using the plotBrowserTrack function and were normalized by the total number of reads in TSS regions^48^. All data were aligned and annotated to the hg38 reference genome.

### Motif enrichment analysis

TF motif enrichment analysis was performed on scATAC peaks using the peakAnnoEnrichment function in ArchR with default parameters based on position frequency matrices from JASPAR 2018 (Supplementary Data 4)^48,55^. Footprinting analysis of TFs was conducted using the getFootprints function in ArchR^48^. To correct for Tn5 insertion bias, footprinting signals were divided by the Tn5 insertion signal prior to plotting.

### SNP selection and LD expansion

Index SNPs implicated in AMD, glaucoma, DR, myopia, or MacTel and located on chromosomes 1-22 and X were collected from the NHGRI-EBI GWAS Catalog, a curated collection of human GWAS^24^. LD expansion was then performed using LDlinkR to add any SNPs in LD with each index SNP^56^, defined as a LD *R*^2^ value >0.9 in the phase 3 genotypes of the 1000 Genomes Project^25^. LD expanded SNPs were filtered to exclude variants in coding regions based on annotations in dbSNP to obtain the final set of noncoding SNPs (Supplementary Data 5)^57^. A list of all GWAS used in this study is provided in Supplementary Table 3.

### HiChIP library generation

H3K27ac HiChIP libraries were prepared as previously reported with minor modifications^14^. Briefly, following isolation of nuclei from frozen retinas as described above, ~8 million nuclei from each sample were washed with nuclei isolation buffer from the diploid chromatin conformation capture (Dip-C) protocol and fixed with 2% paraformaldehyde at room temperature for 10 minutes^58^. Fixed nuclei were then washed twice with cold 1% bovine serum albumin in phosphate-buffered saline before resuspension in 0.5% sodium dodecyl sulfate and resumption of the published HiChIP protocol. Digestion was performed using the MboI restriction enzyme, and sonication was conducted using a Covaris E220 with 5 duty cycles, peak incident power of 140, and 200 cycles per burst for 4 minutes. The ab4729 ChIP validated antibody from Abcam was used to target H3K27ac. HiChIP libraries were sequenced with paired-end 75-bp reads on either an Illumina HiSeq 400 or Illumina NextSeq 550.

### HiChIP data analysis

HiChIP sequencing files were initially processed using the HiC-Pro pipeline (version 2.11.0) to remove duplicate reads, assign reads to MboI restriction fragments, filter for valid interactions, and generate binned interaction matrices^59^. Filtered read pairs from HiC-Pro were subsequently converted into .hic files and inputted into HiCCUPS from the Juicer pipeline to call loops (Supplementary Data 6)^60^. HiChIP interaction maps depicting all valid interactions identified by HiC-Pro were visualized using Juicebox^61^.

### eQTL analysis

Retina eQTL data were obtained from the Eye Genotype Expression (EyeGEx) database^15^. Each of the 1,152 SNPs overlapping with a scATAC peak was searched in the database and the nominal *P* value of any gene associations with that SNP noted. Adjusted *P* values were calculated by multiplying the nominal *P* value by the number of SNP-gene pairs tested for that SNP. Interactions with an adjusted *P* value <0.05 were considered significant.

### Deep learning model training

scATAC-seq reads from the Cell Ranger ARC pipeline were aggregated by cell type to generate cell type-specific fragments files. The fragments files were converted to BigWig tracks of base-resolution Tn5 insertion sites with an +4/-4 shift to account for Tn5 shift. For each cell type, in addition to the peak regions, we selected an equal number of non-peak regions that were matched for GC content in their peaks. We then trained cell type-specific BPNet models to predict the log counts and base-resolution Tn5 insertion profiles as previously reported^8,12^. Briefly, the BPNet model takes as input a 2,114 bp one-hot encoded input sequence and predicts the ATAC-seq profile and log counts in a 1,000 bp window centered at the input sequence. Following BPNet formulation, we used a multinomial negative log likelihood (MNLL) for the profile output of the model and a mean square error (MSE) loss for the log counts output of the model. The relative loss weight used for the counts loss was 0.1 times the mean total counts per region. During each epoch, training examples were jittered by up to 500 bp on either side and a random half of the sequences were reverse complemented. Each batch contained a 10:1 ratio of peaks to non-peak regions. Five models were trained for each cell type corresponding to five disjoint training folds. Model training was performed using Keras/Tensorflow 2. Code used for model training is available at https://github.com/kundajelab/retina-models. Models are available at https://doi.org/10.5281/zenodo.6330053.

### SNP scoring with BPNet

To score LD expanded SNPs associated with eye disease, we centered the input window at the SNP and obtained the log2 fold change in predicted counts between the reference and alternate alleles for each cell type-specific model. We averaged the log2 fold change over the five model folds for each SNP and cell type. To obtain *P* values, we performed one-sided Poisson tests of the predicted alternate allele count with the rate parameter set to the predicted reference allele count (counts averaged over five folds). For each SNP, we combined *P* values across cell types with Fisher’s method and performed Benjamini-Hochberg correction. SNPs with an absolute fold-averaged log2 fold change >0.5 and false discovery rate <0.01 were assigned putative “high effect” annotation. To obtain a background set, random noncoding SNPs were chosen by shuffling a list of all SNPs from the 1000 Genomes Project^25^, filtering out coding regions, and selecting the first 10,000 entries. Only random SNPs localized to chromosomes 1-22 and X were then retained, leaving 9,984 background SNPs. Background SNPs had similar GC content as disease-associated LD expanded SNPs (51% versus 52%) and were scored as described above. Base importance tracks were visualized using Logomaker^62^.

## Data availability

Raw and processed scRNA-seq, scATAC-seq, and HiChIP data from this study have been uploaded to GEO under the accession number GSE196235. A web page summarizing these data is additionally available at https://eyemultiome.su.domains/. BPNet models are available at https://doi.org/10.5281/zenodo.6330053. All data generated in this study are available upon reasonable request.

## Code availability

Code used for BPNet model training is available at https://github.com/kundajelab/retina-models. All other custom code used in this study is available upon request.

## ACKNOWLEDGEMENTS

The authors are grateful to Lions VisionGift, Lions Gift of Sight, and the donors who made this study possible. Computing for this project was performed on the Sherlock cluster, a resource provided and maintained by the Stanford Research Computing Center. This work was supported by National Institutes of Health (NIH) grant RM1-HG007735 (to H.Y.C.). S.K.W. is supported by a National Eye Institute training grant (T32EY027816). H.Y.C. is an Investigator of the Howard Hughes Medical Institute.

## AUTHOR CONTRIBUTIONS

S.K.W. and H.Y.C. conceived the project and designed experiments. S.K.W., R.L., K.K., and C.L. performed experiments. S.K.W., S.N., A.P., A.P., and J.B.K. performed data analysis. A.K. and H.Y.C. supervised the work. S.K.W. and H.Y.C. wrote the manuscript with input from all authors.

---

**Extended Data Figure 1.**
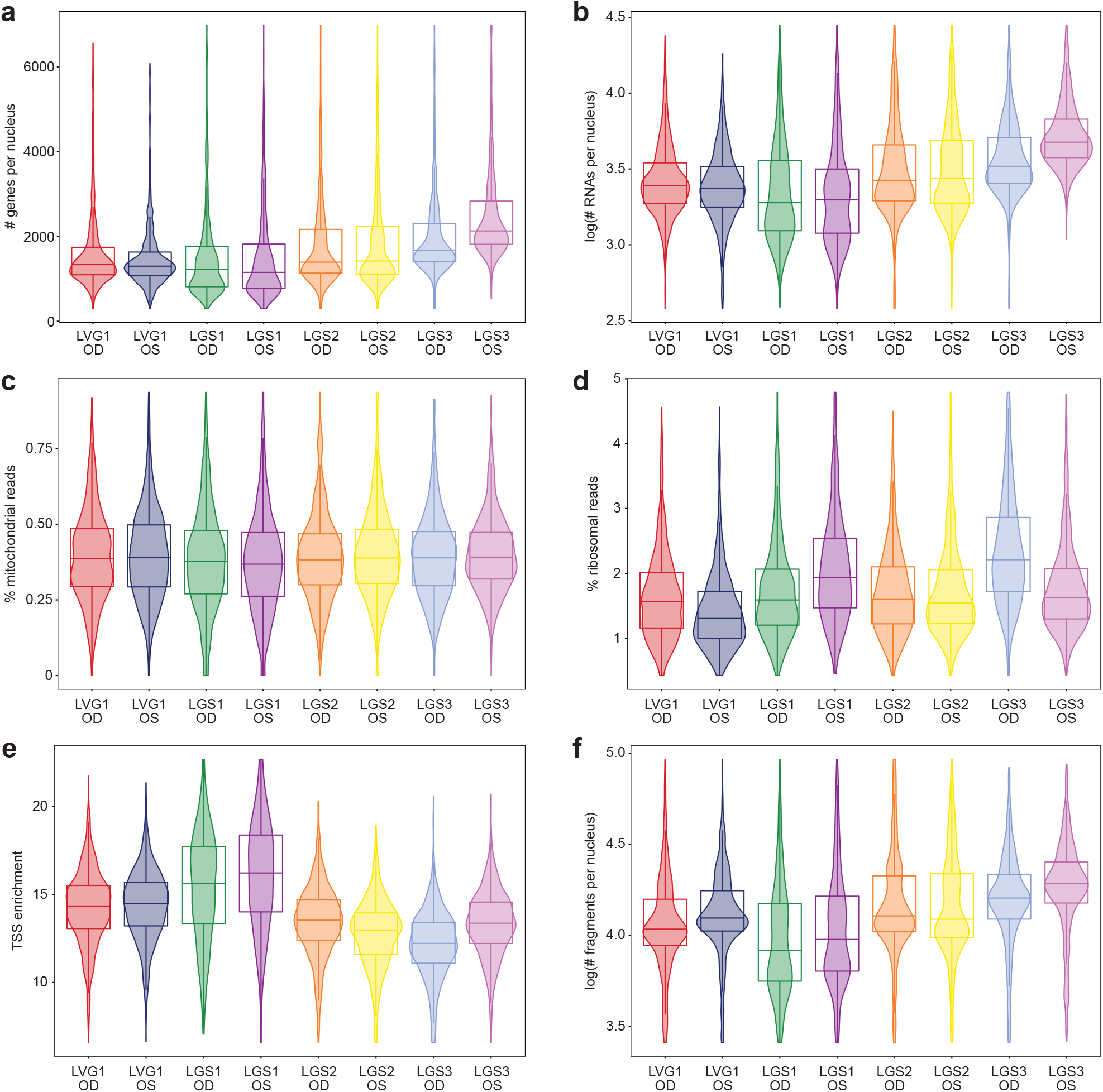
Single-cell RNA- and ATAC-seq quality control metrics. **a-f**, Violin plots depicting the number of detected genes per nucleus (a), number of detected RNA transcripts per nucleus (b), percentage of mitochondrial reads (c), percentage of ribosomal reads (d), TSS enrichment (e), and number of detected fragments per nucleus (f) by retina in the final dataset. Boxes depict the 25^th^ percentile, median, and 75^th^ percentile of each metric.

**Extended Data Figure 2.**
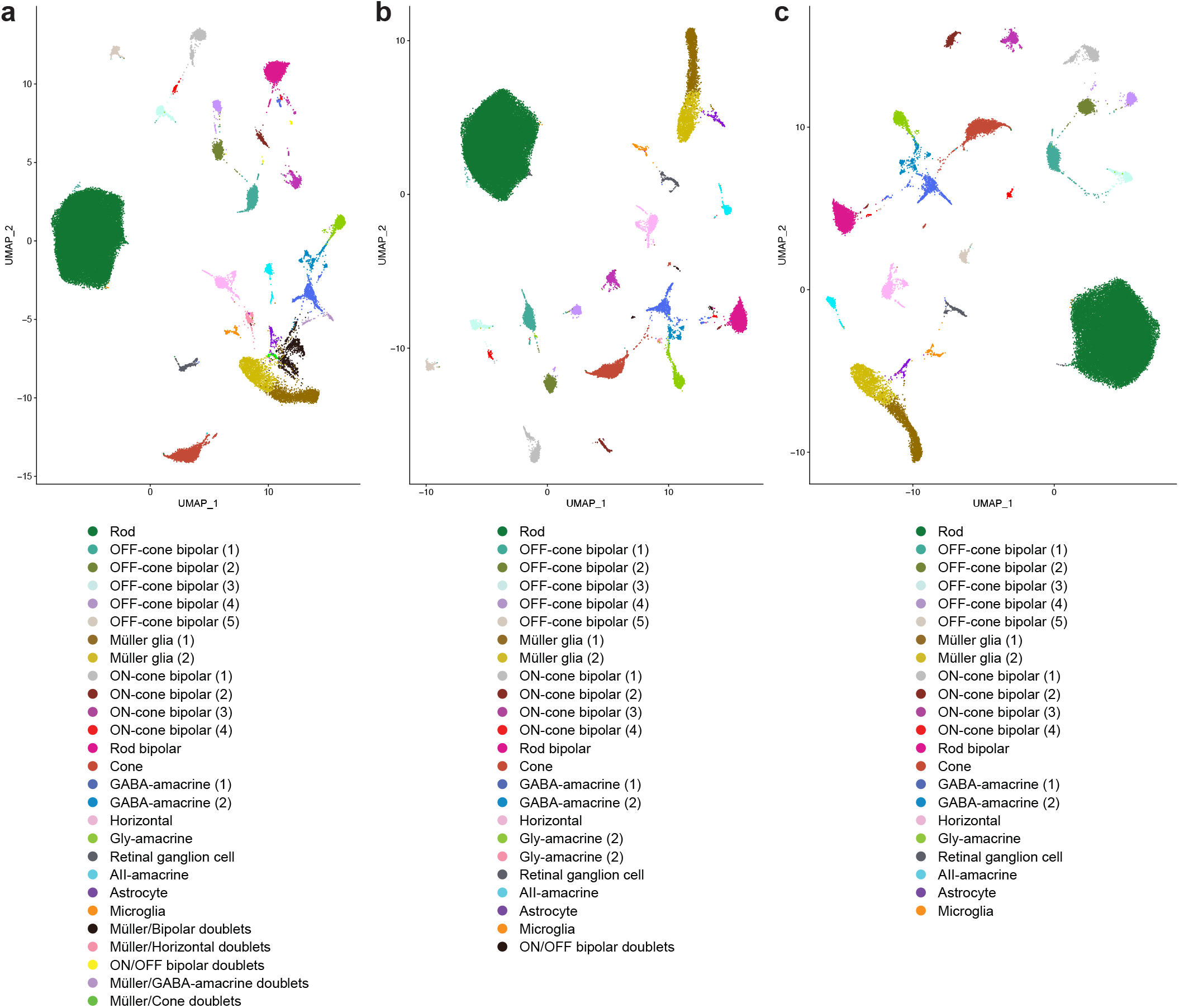
Single-cell RNA-seq cluster assignments. **a**, First iteration uniform manifold approximation and projection (UMAP) plot of the scRNA-seq dataset after quality control filtering, automated removal of doublets, and exclusion of clusters with no detected marker genes. Five clusters comprised of putative doublets were subsequently removed and the dataset re-clustered. **b**, Second iteration UMAP of the scRNA-seq dataset. One cluster comprised of putative doublets was subsequently removed and the dataset re-clustered. **c**, Final iteration UMAP of the scRNA-seq dataset. Clusters representing subpopulations of the same cell type were grouped together for downstream analyses.

**Extended Data Figure 3.**
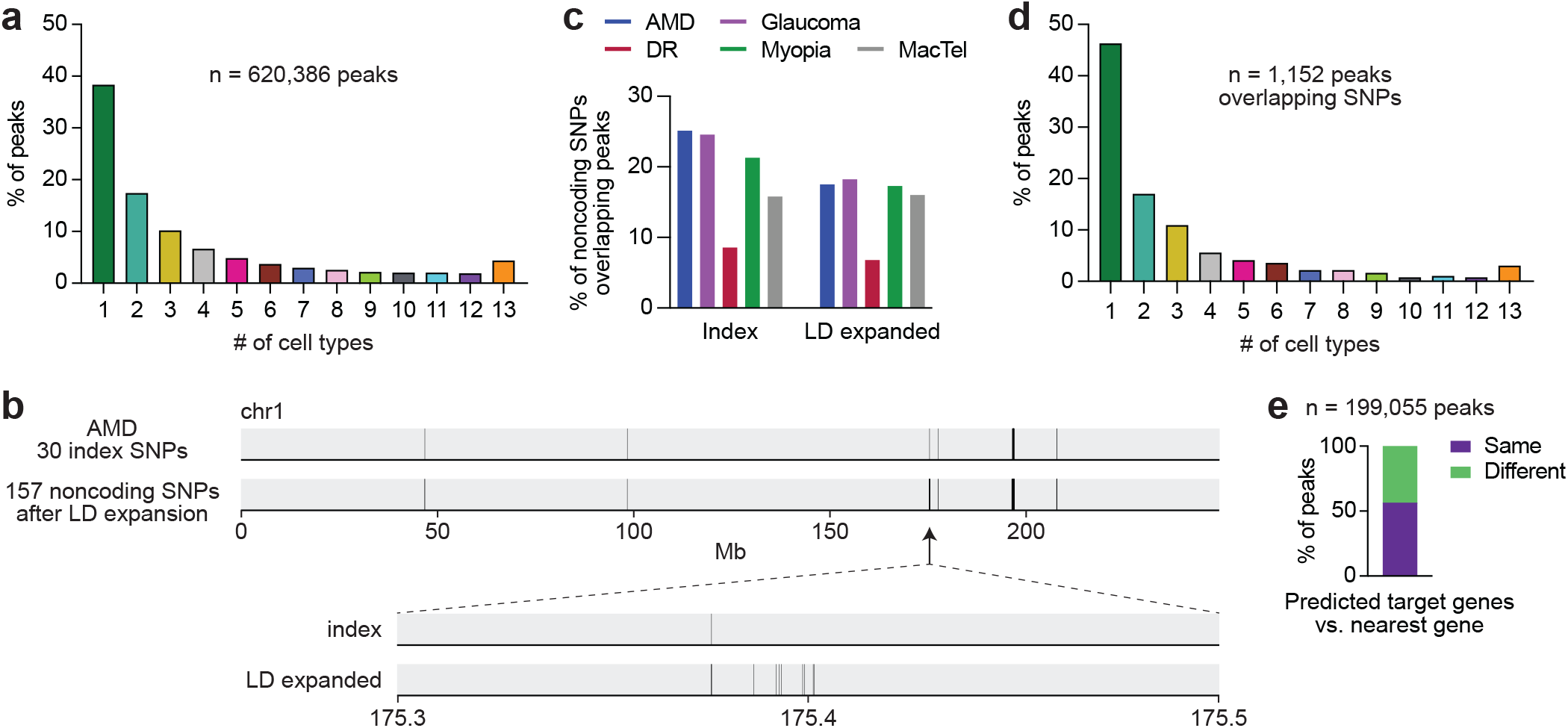
Characterization of chromatin accessibility peaks and linkage disequilibrium expanded SNPs. **a**, Number of cell types exhibiting each of the 620,386 scATAC peaks. **b**, Visual depiction of LD expansion for the 30 index SNPs on chromosome 1 associated with AMD. Each vertical black line represents a SNP. **c**, Percentage of noncoding index and LD expanded SNPs from each disease that overlapped with at least one scATAC peak. **d**, Number of cell types exhibiting each of the 1,152 scATAC peaks that overlapped with a LD expanded SNP. **e**, Comparison of predicted target genes with the nearest gene for each chromatin accessibility peak with at least one predicted target gene. Same was defined as any of the predicted target genes being the nearest gene.

**Extended Data Figure 4.**
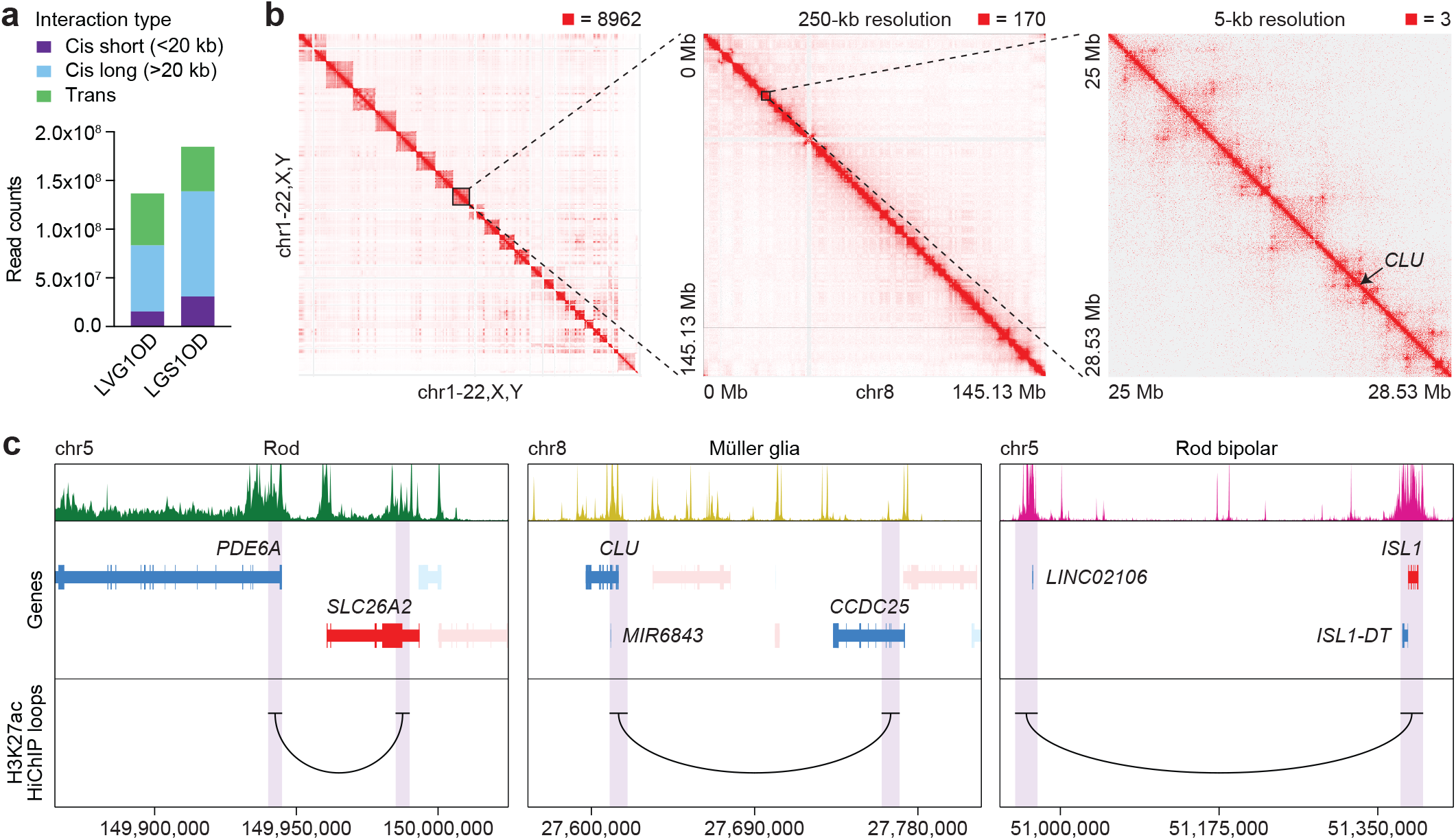
H3K27ac HiChIP predicts enhancer-promoter interactions in the human retina. **a**, HiChIP interaction types by sample. Cis refers to interactions on the same chromosome, while trans refers to interactions spanning separate chromosomes. **b**, HiChIP interaction maps at whole genome, 250-kb, and 5-kb resolution. Sample shown is LGS1OD. Signal was normalized to the square root of coverage. Numbers above the interaction maps indicate maximum signal in each matrix. **c**, Sequencing tracks of cell type chromatin accessibility and H3K27ac HiChIP loops overlapping with the TSS of selected marker genes. Genes in the sense and antisense directions are shown in red and blue, respectively. Regions encompassed by the loop anchors are highlighted in purple.

**Extended Data Figure 5.**
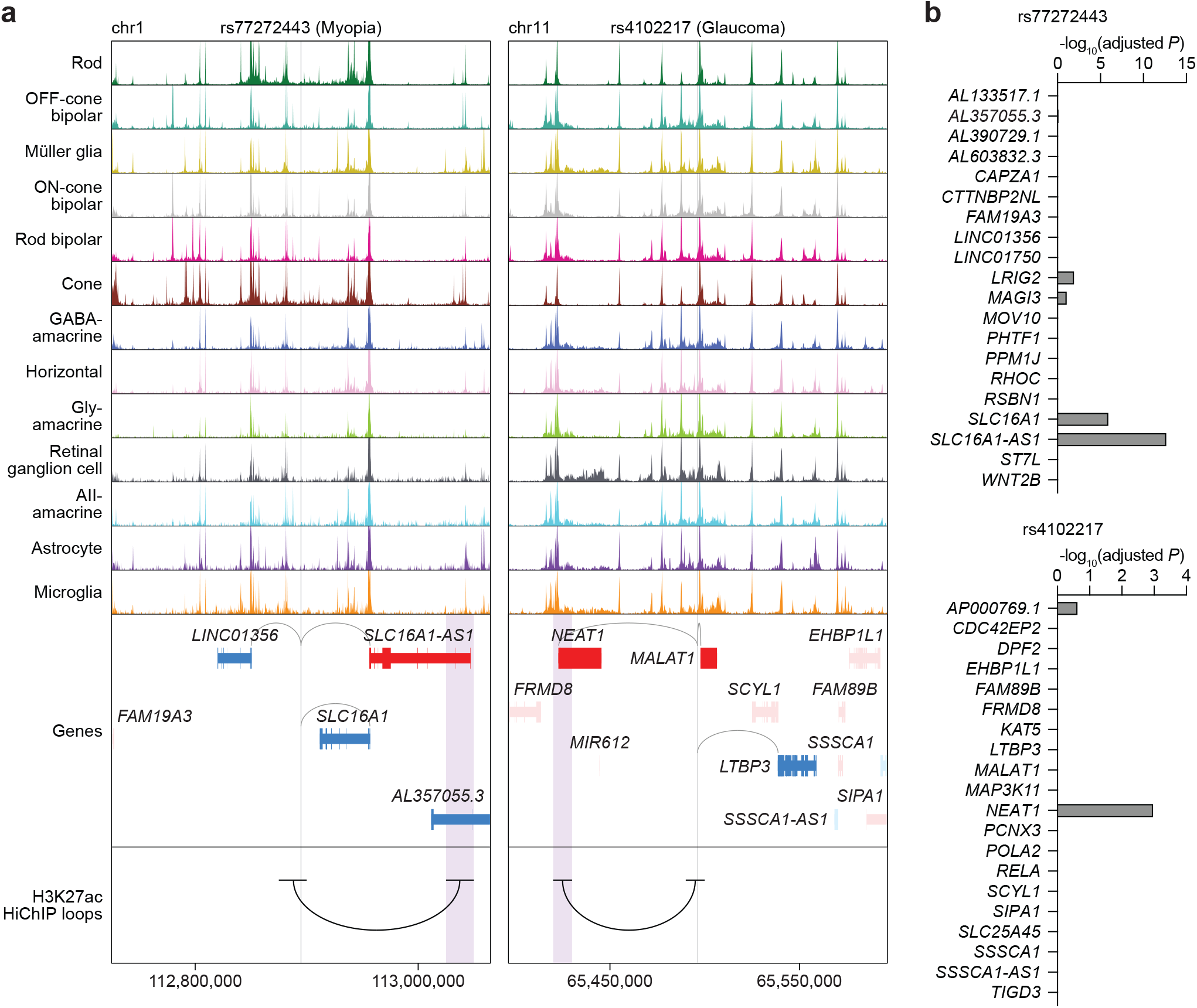
Integration of single-cell multiome with HiChIP and eQTL data refines SNP-target gene predictions. **a**, Sequencing tracks of chromatin accessibility near rs77272443 (chr1:112894884) and rs4102217 (chr11:65496424). Genes in the sense and antisense directions are shown in red and blue, respectively. The location of each SNP is depicted by a vertical gray line. Gray arcs indicate predicted target genes for the scATAC peak containing the SNP of interest. Black arcs overlapping with SNPs indicate H3K27ac HiChIP loops with the regions encompassed by the opposite anchors highlighted in purple. **b**, Significance of SNP-gene associations for rs77272443 or rs4102217 and their 20 nearest genes as determined by retina eQTL analysis. Adjusted *P* values for each gene were calculated by multiplying the nominal *P* value listed in the EyeGEx database by the number of SNP-gene pairs tested for that SNP.

**Extended Data Figure 6.**
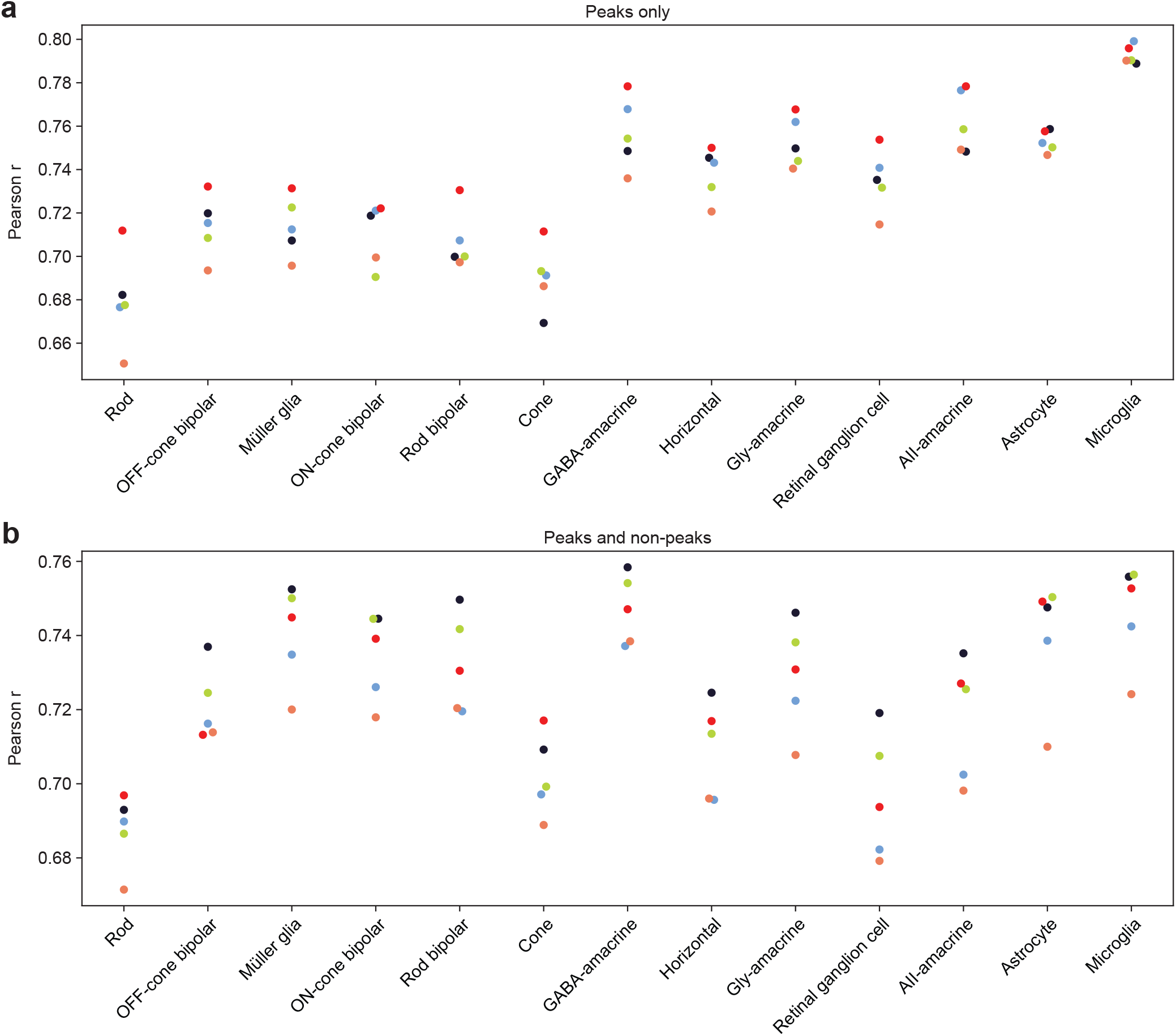
BPNet model performance. **a,b**, Correlation scores between predicted and actual observed log counts in only peak (a) or both peak and non-peak (b) regions on chromosome sequences that were withheld during cell typespecific model training. Each color represents one of five model folds.

**Extended Data Figure 7.**
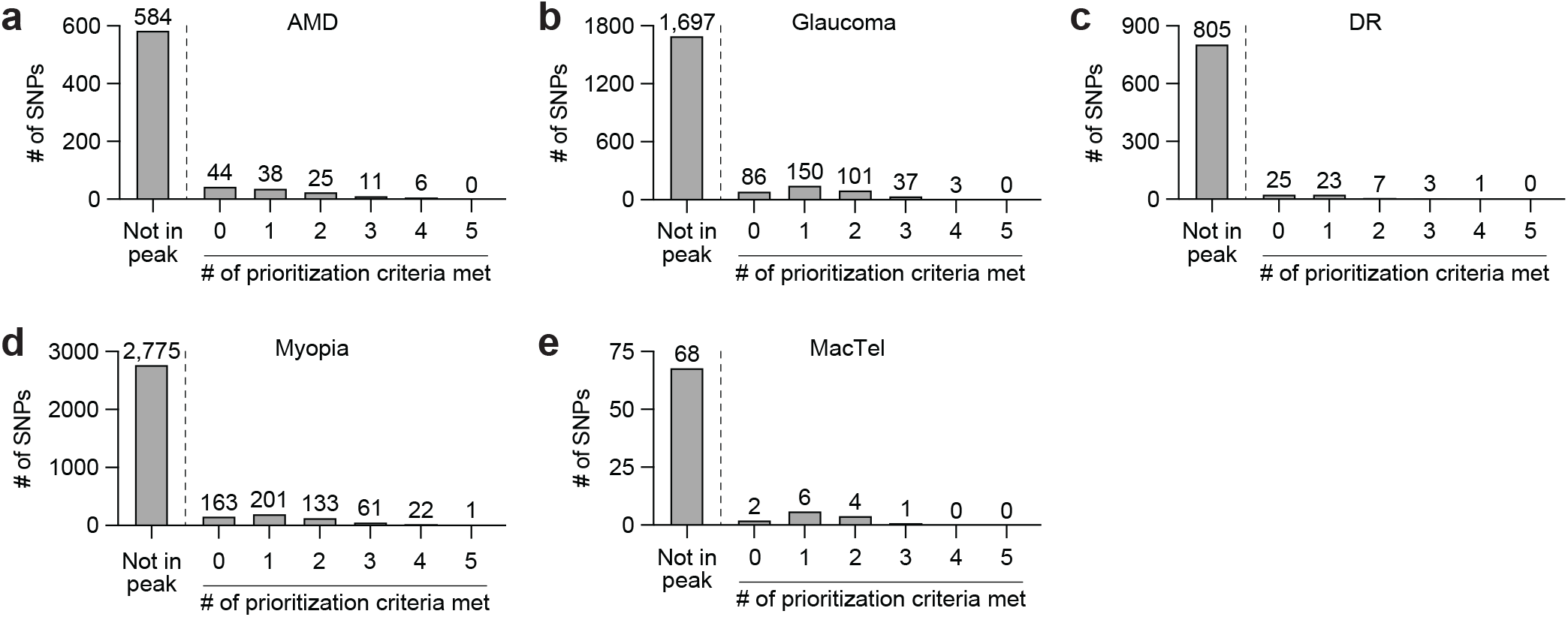
Summary of SNP prioritization. **a-e**, Total number of prioritization criteria met by LD expanded SNPs in loci associated with AMD (a), glaucoma (b), DR (c), myopia (d), and MacTel (e). Prioritization criteria were 1) coaccessibility with a promoter peak, 2) accessibility correlated with expression of a nearby gene, 3) linkage to a gene by H3K27ac HiChIP, 4) significant association with a gene by retina eQTL data, and 5) high effect designation by deep learning.

