## Supplementary Tables and Information for "Single-cell multiome of the human retina and deep learning nominate causal variants in complex eye diseases"

**Supplementary Table 1. Donor information**

| ID | Age | Sex | Death-to-preservation interval | Cause of death |
| --- | --- | --- | --- | --- |
| LVG1 | 55 | M | 9 hours | Cardiac arrest |
| LGS1 | 85 | F | 11 hours | Sepsis |
| LGS2 | 87 | M | 9 hours | Fall |
| LGS3 | 74 | F | 12 hours | Dementia |

**Supplementary Table 2. Cell counts per retina**

| Cell type | LVG1  OD | LVG1  OS | LGS1  OD | LGS1  OS | LGS2  OD | LGS2  OS | LGS3  OD | LGS3  OD |
| --- | --- | --- | --- | --- | --- | --- | --- | --- |
| Rod | 3151 | 3061 | 6132 | 6777 | 3409 | 3190 | 3299 | 3040 |
| OFF-cone bipolar | 398 | 373 | 447 | 558 | 557 | 622 | 549 | 432 |
| Müller glia | 384 | 372 | 496 | 448 | 608 | 559 | 436 | 367 |
| ON-cone bipolar | 223 | 245 | 299 | 327 | 449 | 393 | 373 | 308 |
| Rod bipolar | 234 | 190 | 358 | 263 | 350 | 302 | 313 | 314 |
| Cone | 197 | 164 | 335 | 386 | 325 | 365 | 240 | 211 |
| GABA-amacrine | 194 | 135 | 257 | 257 | 169 | 240 | 171 | 120 |
| Horizontal | 116 | 107 | 114 | 129 | 194 | 124 | 174 | 177 |
| Gly-amacrine | 83 | 66 | 114 | 138 | 77 | 106 | 80 | 66 |
| Retinal ganglion cell | 98 | 54 | 26 | 20 | 116 | 53 | 100 | 26 |
| AII-amacrine | 42 | 53 | 94 | 72 | 41 | 83 | 43 | 23 |
| Astrocyte | 40 | 35 | 4 | 8 | 31 | 19 | 59 | 36 |
| Microglia | 15 | 13 | 33 | 22 | 39 | 51 | 29 | 30 |
| Total | 5175 | 4868 | 8709 | 9405 | 6365 | 6107 | 5866 | 5150 |

**Supplementary Table 3. First author and year of GWAS by disease**

| AMD | Glaucoma | DR | Myopia | MacTel |
| --- | --- | --- | --- | --- |
| Klein 2005^1^ | Thorleifsson 2007^2^ | Huang 2011^3^ | Nakanishi 2009^4^ | Scerri 2017^5^ |
| Chen 2010^6^ | Meguro 2010^7^ | Grassi 2011^8^ | Li 2011^9^ | Bonelli 2021^10^ |
| Neale 2010^11^ | Burdon 2011^12^ | Sheu 2013^13^ | Li 2011^14^ |  |
| Kopplin 2010^15^ | Nakano 2012^16^ | Awata 2014^17^ | Shi 2011^18^ |  |
| Yu 2011^19^ | Osman 2012^20^ | Burdon 2015^21^ | Fan 2012^22^ |  |
| Arakawa 2011^23^ | Wiggs 2012^24^ | Graham 2018^25^ | Meng 2012^26^ |  |
| Cipriani 2012^27^ | Takamoto 2012^28^ | Meng 2018^29^ | Shi 2013^30^ |  |
| Sobrin 2012^31^ | Vithana 2012^32^ | Pollack 2019^33^ | Khor 2013^34^ |  |
| Holliday 2013^35^ | Nakano 2014^36^ | Meng 2019^37^ | Simpson 2014^38^ |  |
| Fritsche 2013^39^ | Hoffmann 2014^40^ | Liu 2019^41^ | Pickrell 2016^42^ |  |
| Naj 2013^43^ | Chen 2014^44^ | Hsieh 2020^45^ | Tedja 2018^46^ |  |
| Cheng 2015^47^ | Gharahkhani 2014^48^ | Imamura 2021^49^ | Boutin 2020^50^ |  |
| Fritsche 2016^51^ | Aung 2015^52^ |  | Meguro 2020^53^ |  |
| Ruamviboonsuk 2017^54^ | Li 2015^55^ |  | Tideman 2021^56^ |  |
| Persad 2017^57^ | Vishal 2016^58^ |  |  |  |
| Yan 2018^59^ | Khor 2016^60^ |  |  |  |
| Han 2020^61^ | Verma 2016^62^ |  |  |  |
| Winkler 2020^63^ | Bailey 2016^64^ |  |  |  |
| Guenther 2020^65^ | Zagajewska 2018^66^ |  |  |  |
|  | Gharahkhani 2018^67^ |  |  |  |
|  | Choquet 2018^68^ |  |  |  |
|  | MacGregor 2018^69^ |  |  |  |
|  | Bonnemaijer 2018^70^ |  |  |  |
|  | Zhou 2018^71^ |  |  |  |
|  | Shiga 2018^72^ |  |  |  |
|  | Hauser 2019^73^ |  |  |  |
|  | Craig 2020^74^ |  |  |  |
|  | Kim 2020^75^ |  |  |  |
|  | Ishigaki 2020^76^ |  |  |  |
|  | Gharahkhani 2021^77^ |  |  |  |
|  | Sakaue 2021^78^ |  |  |  |

**Supplementary Data 1**

Marker genes from scRNA-seq by cell type

**Supplementary Data 2**

Bed format of reproducible scATAC peaks by cell type

**Supplementary Data 3**

Bed format of scATAC marker peaks and nearest genes by cell type

**Supplementary Data 4**

-log10 adjusted *P* values for TF motifs by cell type

**Supplementary Data 5**

Summary of noncoding SNPs and prioritization results

**Supplementary Data 6**

H3K27ac HiChIP loops and intersecting genes

Supplementary Data 7

BPNet scores for disease-associated and randomly selected SNPs by cell type
